# First cleavage is a manifestation of the geometry of the unfertilized oocyte: implications for monozygotic twinning in mice

**DOI:** 10.1101/2023.12.30.572752

**Authors:** Thomas Nolte, Reza Halabian, Steffen Israel, Yutaka Suzuki, Georg Fuellen, Wojtek Makalowski, Michele Boiani

**Affiliations:** Max Planck Institute for Molecular Biomedicine, Röntgenstrasse 20, 48149 Muenster, Germany; Faculty of Medicine, Institute of Bioinformatics, University of Münster, 48149 Münster, Germany; Department of Computational Biology and Medical Sciences, Graduate School of Frontier Sciences, University of Tokyo, Chiba 277-8562, Japan; Rostock University Medical Center, Institute for Biostatistics and Informatics in Medicine and Aging Research (IBIMA), Ernst-Heydemann-Strasse 8, 18057 Rostock, Germany

## Abstract

A long-standing question in mammalian embryology is whether regional differences of oocyte composition matter for the properties of blastomeres receiving those regions after fertilization. A hitherto untested hypothesis is that allocation depends on the orientation of 1^st^ cleavage. However, the orientation is influenced by the site of sperm entry, which can be almost anywhere on the membrane of oocytes when these are inseminated. This variability undermines consistency and reproducibility of studies. Therefore, we harnessed the intracytoplasmic sperm injection to impose the site of fertilization in three specific ooplasmic regions (animal pole, vegetal pole, equator) in mice. Notwithstanding this categorical distinction, after 1^st^ cleavage, the sister blastomeres differed from each other nearly the same way, as measured by gene expression and twin blastocysts formation following 2-cell embryo splitting. We reasoned that either the oocyte territories did not matter, or their effect was obscured by other factors. To shed light on these possibilities, we immobilized the oocytes on the micromanipulation stage during sperm injection and for 24 h thereafter. Imaging revealed that the orientation of 1^st^ cleavage, instead of varying with the fertilization site, followed the shorter diameter of the unfertilized oocyte. This led in most cases to the segregation of animal and vegetal hemispheres into the sister blastomeres of 2-cell embryos. Since one blastomere received more of the animal materials and the other blastomere more of the vegetal materials, this offers a rationale to explain the distinct properties of monozygotic twins derived from 2-cell embryos in mice.

## Introduction

For more than sixty years scholarly books of developmental biology taught that the 2-cell stage blastomeres in mammals were as totipotent as the zygote, that is, they had same ability to produce an entire individual after dissociation into single cells (Tarkowski, 1959). This notion was fostered by the historical studies of Hans Driesch performed in 2-cell sea urchin embryos at the end of the 19^th^ century (Driesch, 1964 (1892)). Each dissociated blastomere would develop into a pluteus larva, albeit not always of the same size, and this behavior seemed to relate to how the plane of 1^st^ cleavage was oriented. From 2017 onwards, increasing evidence has been showing that when mouse blastomeres are separated at the 2-cell stage, only one blastomere is able to develop into a viable blastocyst or develop fully into a live mouse in most cases (Casser *et al*., 2017; Krawczyk *et al*., 2021; Maemura *et al*., 2021). Full developmental potential is probably attained when the blastomere progeny accrues sufficient epiblast cells by the blastocyst stage (Morris *et al*., 2012). In addition to these functional features, also molecular differences have been identified between mouse blastomeres at the 2-cell stage (Hupalowska *et al*., 2018; Wang *et al*., 2018; Wang *et al*., 2021). These findings triggered a re-assessment of the notion that 2-cell blastomeres have identical developmental potential (Boiani *et al*., 2019). Then again, how the differences originate remains largely unknown.

Recently, we rediscovered studies written in German between the early 1950s and the early 1960s, which proposed that inter-blastomere differences depend, at least in part, on the partition of oocyte territories through the orientation of the 1^st^ cleavage after fertilization in rabbits (Seidel, 1952; Seidel, 1960). Subsequent studies in mice proposed that this orientation was related to the topology of fertilization e.g. to where the spermatozoon had entered the ooplasm and how the two pronuclei apposed to each other (Hiiragi and Solter, 2004; Piotrowska and Zernicka-Goetz, 2001; Piotrowska-Nitsche and Chan, 2013). These two proposals and their consequences have largely remained untested, although they could help explain why the sister blastomeres in 2-cell mouse embryos are different from each other (Casser *et al*., 2017). An obvious difficulty in testing the two proposals is that the sperm entry point can be located almost anywhere on the membrane (oolemma) of inseminated oocytes, and that mammalian oocytes lack obvious territories that can be discerned in the living state, except for the animal and vegetal poles. The animal pole is landmarked by the 1^st^ polar body lying on top of the meiotic spindle (Solter, 2016), whereby the two structures lie on the same side (ipsilateral). Diametrically opposite the animal pole is the vegetal pole (contralateral), which in mice contains a higher density of endoplasmic reticulum not visible to the naked eye (Kline *et al*., 1999). The two poles define an animal-vegetal (A-V) axis, which coincides with the future plane of 1^st^ zygotic division called ‘meridional’ according to some investigators (Ajduk and Zernicka-Goetz, 2016), while according to other investigators it does not (Hiiragi and Solter, 2004). The situation might be more complex, as it was reported that division perpendicular to the A-V axis (i.e. equatorial) occurred in a non-negligible proportion of mouse zygotes (Gardner and Davies, 2003). These different topologies are envisioned to have opposite consequences: if the cleavage axis was meridional i.e. along the A-V axis, then the two blastomeres should inherit equal proportions of animal and vegetal materials, and the cellular properties could be similar to each other if not identical; while if the cleavage axis was equatorial i.e. perpendicular to the A-V axis, then the two blastomeres should inherit different proportions of those materials, and the cellular properties could be distinct. We use the conditional tense because the molecular nature of such materials and the boundary between them has been a matter of debate for many years. Whole-mount immunohistochemistry studies pointed at molecular candidates (e.g. leptin, STAT3, TGFβ2, VEGF, BCL-X, BAX) that occupied polarized domains in mouse and human oocytes (Antczak and Van Blerkom, 1997; 1999; Schulz and Roberts, 2011). However, when immunohistochemistry was conducted on histological sections instead of whole-mounts, the polarized domains were not as pronounced (Littwin and Denker, 2011). As to the boundary, prevalent view is that it is perpendicular to the A-V axis, however, historical histochemical studies by Albert Dalcq suggested that the boundary could be parallel with rather than perpendicular to the A-V axis (Dalcq, 1957).

In this study, we sought to get an experimental grip on a problem that so far impeded to ascertain if the transmission of animal and vegetal materials factors in to shape the properties of 2-cell stage blastomeres, namely: the natural variability in the site of sperm penetration on the oolemma. To this end, we harnessed the method of intracytoplasmic sperm injection in mice (ICSI; (Kimura and Yanagimachi, 1995)). We applied ICSI at the animal pole or vegetal pole or half way between the poles (equatorial region) of metaphase II (MII) oocytes, which we called “site-specific ICSI”. We ascertained that all three fertilization topologies were conducive to birth, even the one at the animal pole, which is considered detrimental (Blake *et al*., 2000; Labs *et al*., 2016; Mori *et al*., 2021; Plusa *et al*., 2002). After cleavage, we dissociated the 2-cell stage blastomeres and subjected them to either molecular or functional analysis, expecting to find that the properties of the sister blastomeres varied with the site of fertilization. Contrary to expectation, the differences between the sister blastomeres were almost the same no matter the three fertilization sites. This suggested the following: either the oocyte territories did not matter, or their effect was evened out by other factors. To shed light on these possibilities, we immobilized the oocytes on the micromanipulation stage for 24 h to see how the oocyte divided after site-specific ICSI. This approach revealed that majority of oocytes divided quasi-equatorially with respect to their cell geometry prior to fertilization, apparently following the Hertwig’s rule of the shorter cell diameter (Hertwig, 1893), irrespective of the fertilization site. This means that one blastomere receives more of the animal materials and the other more of the vegetal materials, thus offering a rationale for the differences between monozygotic twins produced via blastomere separation at the 2-cell stage (Casser *et al*., 2017).

## Results

### Site-specific ICSI allows for standardized sperm deposition at the poles or equator of MII mouse oocytes and supports full development

Using ICSI as method of fertilization and the meiotic spindle as a stationary marker to identify the oocyte’s animal pole, we placed the sperm head at one of only three specific and reproducible locations, thereby reducing the variability of the fertilization site that is otherwise encountered when oocytes are fertilized *in vivo* or inseminated *in vitro*. For more consistency, we used throughout the study the same batch of spermatozoa, which were aliquoted and cryopreserved for the purpose.

Using a micromanipulator (Figure 1A) we performed ICSI into MII mouse oocytes (Figure 1B) that were oriented so as to have the meiotic spindle sitting at 3, 9 or 12 o’clock when introducing single sperm heads always from the 9 o’clock position toward the 3 o’clock position (Figure 1C,D,E). As a result, seen from the standpoint of the spindle, these oocytes became fertilized ipsilaterally (animal pole), contralaterally (vegetal pole) or half-way between (equatorially). Immediate oocyte survival rates after ICSI were similar across the three groups: ipsilateral 81 ± 8% (566 injections in 14 sessions), contralateral 85 ± 9% (608 injections in 14 sessions) and equatorial 87 ± 6% (626 injections in 14 sessions). When removed from the micromanipulator, cultured in a regular incubator and then briefly returned to the micromanipulator for imaging 5 hours later, the fertilized oocytes had invariably formed 2 pronuclei (Figure 1C’,D’,E’), notwithstanding a concern that the sperm nucleus could be dragged along with the 2^nd^ polar body if placed too close to the spindle (Mori *et al*., 2021).

**Figure 1.**
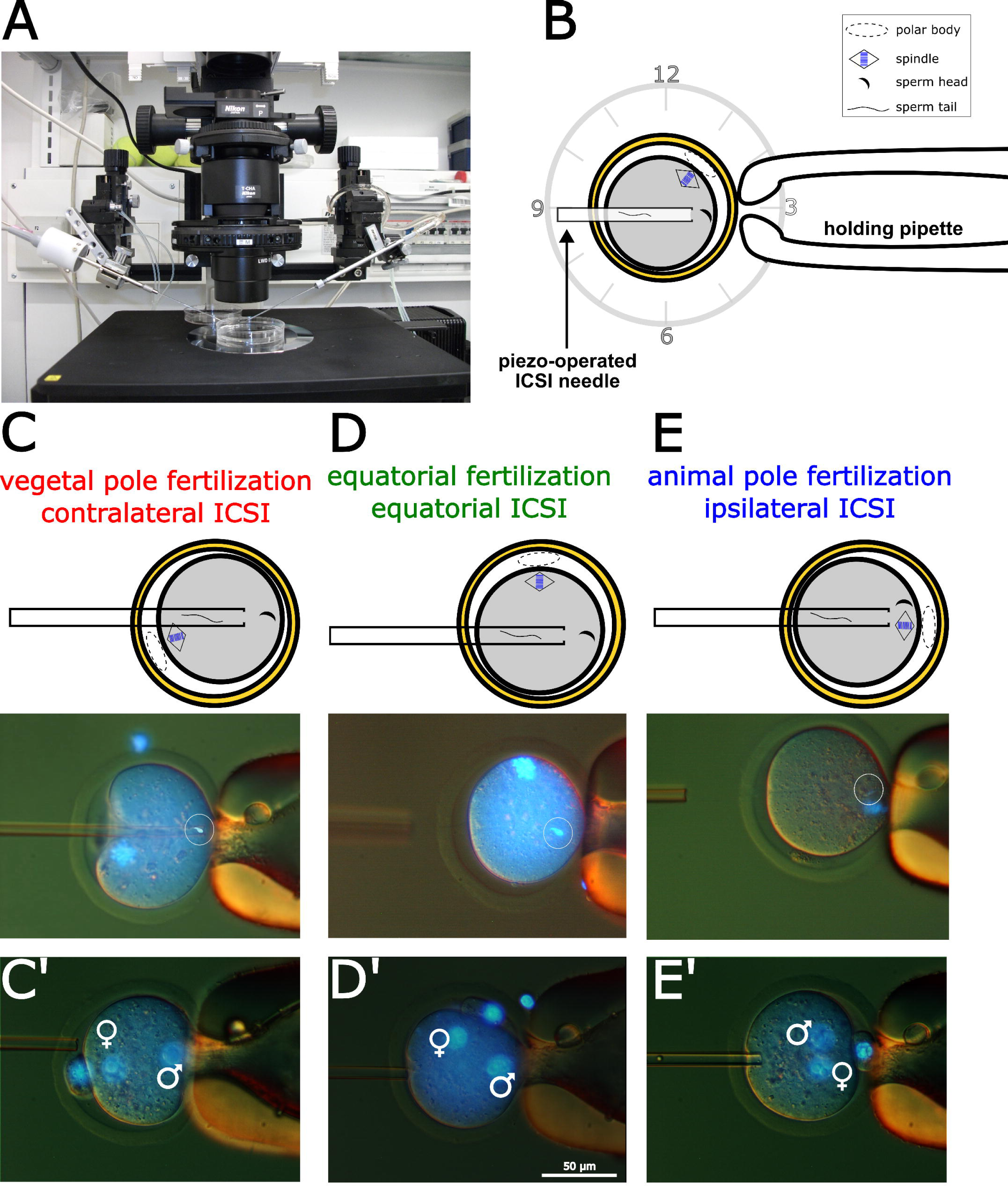
The site-specific ICSI method used in this study to standardize the fertilization topology in respect to presumptive oocyte territories. Stage of the micromanipulator (A) on which the ICSI (B) was performed. (B) MIi mouse oocytes (B6C3Fl) were held in position by suction of the zona pellucida (ZP) with a holding pipette at the 3 o’clock position, and injected with a single sperm head (CDl) using a piezo-driven needle approaching from the 9 o’clock position. C, D, E: a single sperm head (encircled) was deposited in the ooplasm opposite the spindle i.e. contralateral (vegetal pole), next to the spindle i.e. ipsilateral (animal pole) or half-way in between (equator). C’, D’, E’: two pronuclei (d’9) had formed 5 hours after ICSI, attesting that the activation of the oocytes had been successful.

We ascertained that the site-specific ICSI had no bold negative effects due to local perturbations, given a widespread assumption that the animal pole is a sensitive location that’s best not to perturb (Labs *et al*., 2016; Plusa *et al*., 2002). To ascertain this, we allowed for further development. Cultured four days inside an incubator, the pronuclear oocytes progressed to blastocysts irrespective of the ICSI site, with a significant reduction in blastocyst rate (Figure 2A) but not in total cell number (Figure 2B) relative to control embryos formed by oocytes fertilized *in vivo* and cultured *in vitro* (naturally fertilized). Following surgical transfer of the blastocysts into foster uteri of pseudo-pregnant mice, birth rates were reduced – albeit not significantly - in the ICSI groups relative to control group, particularly in the ipsilateral ICSI group (Figure 2C). Presumably this generalized reduction of viability was the tradeoff for the higher consistency of using the same batch of cryopreserved spermatozoa throughout the study, as it is well known that cryopreservation is accompanied by cryopreservation-associated damage.

**Figure 2.**
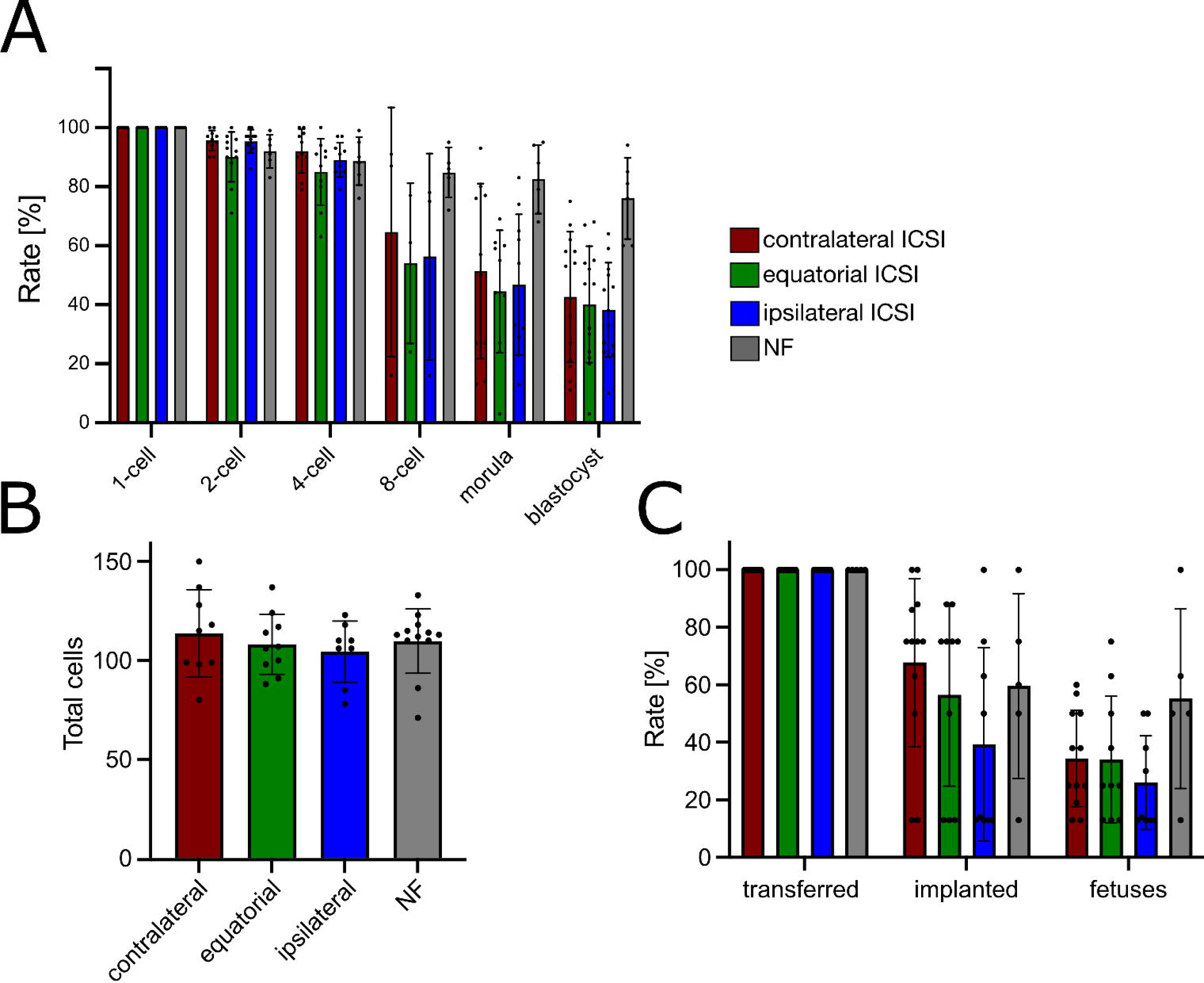
Validation of the site-specific ICSI method by way of subsequent development. **(A)** Blastocysts formed at rates not significantly different from each other in all three ICSI groups (contralateral 43 ± 22%, equatorial 40 ± 20%; ipsilateral 38 ± 16%; p > 0.555; Wilcoxon test), albeit lower than those of oocytes subjected to natural fertilization (NF) (76 ± 14%; p < 0.004; Wilcoxon test). Starting N of zygotes in (A): contralateral = 464; equatorial = 512; ipsilateral = 396; NF= 779. **(B)** Blastocysts’ total cell numbers were similar across all groups (contralateral ICSI 114 ± 6; equatorial ICSI 108 ± 6; ipsilateral ICSI 104 ± 6; NF 110 ± 5; p > 0.229; Wilcoxon test). N of blastocysts counted in (B): contralateral = 9; equatorial = 10; ipsilateral = 8; NF = 12. **(C)** After transfer to the uteri of pseudo-pregnant females, blastocysts from oocytes fertilized by ICSI developed to term at rates not significantly different from each other (contralateral =34 ± 17%; equatorial= 34 ± 22%; ipsilateral = 26 ± 16%; p > 0.233, Wilcoxon test) albeit lower than the rates of NF counterparts (55 ± 31%; p > 0.073, Wilcoxon test). Starting N of transferred blastocysts and recipient mothers, respectively, used in (C): contralateral ICSI = 92, 12; equatorial ICSI =72, 10; ipsilateral ICSI = 73, 9; NF= 40, 5. Data in **(A)(B)(C)** are presented as means and standard deviations; black dots represent the individual replicates.

The results of this section can be summarized as follows. Site-specific ICSI allowed for standardization of zygote production with respect to fertilization topology. All three fertilization sites (including the animal pole of the ipsilateral ICSI) were conducive to birth. These outcomes support that the events observed here *in vitro* reflected, to a reasonable extent, the natural (*in vivo*) events.

### Despite three different sites of oocyte fertilization, resultant blastomeres behave the same way in terms of gene expression

We hypothesized that if ICSI at the animal or vegetal pole (ipsilateral, contralateral) or at the equator resulted in systematic partitioning oocyte territories into the blastomeres at 1^st^ cleavage, then it should leave a footprint on the molecular composition of the blastomeres, and each of the three fertilization topologies should have its own footprint. It should be feasible to discern the footprints if a decent number of blastomere pairs was analyzed. To this end, 50 pairs of blastomeres from each ICSI group (150 pairs in total) were extracted from the zona pellucida (ZP; Figure 3A) as per our established method (Casser *et al*., 2017), and subjected to single-cell transcriptome profiling by RNA sequencing (RNA-seq). With an aim to minimize batch effects and day-to-day variation, we adopted the following four measures for the purpose RNA-seq: 1) the contralateral, equatorial and ipsilateral ICSIs were performed on the same day, 2) the 2-cell embryos were bisected on the same day, 3) the individual blastomeres were transferred to lysis buffer in a Mosquito® plate only if the 2^nd^ polar body had remained trapped inside the evacuated ZP, and 4) the RNA extraction was conducted in parallel for all samples. Following RNA-seq the average sequencing depth was 3.5 ± 1.3 million uniquely mapped reads per specimen of single blastomeres.

**Figure 3.**
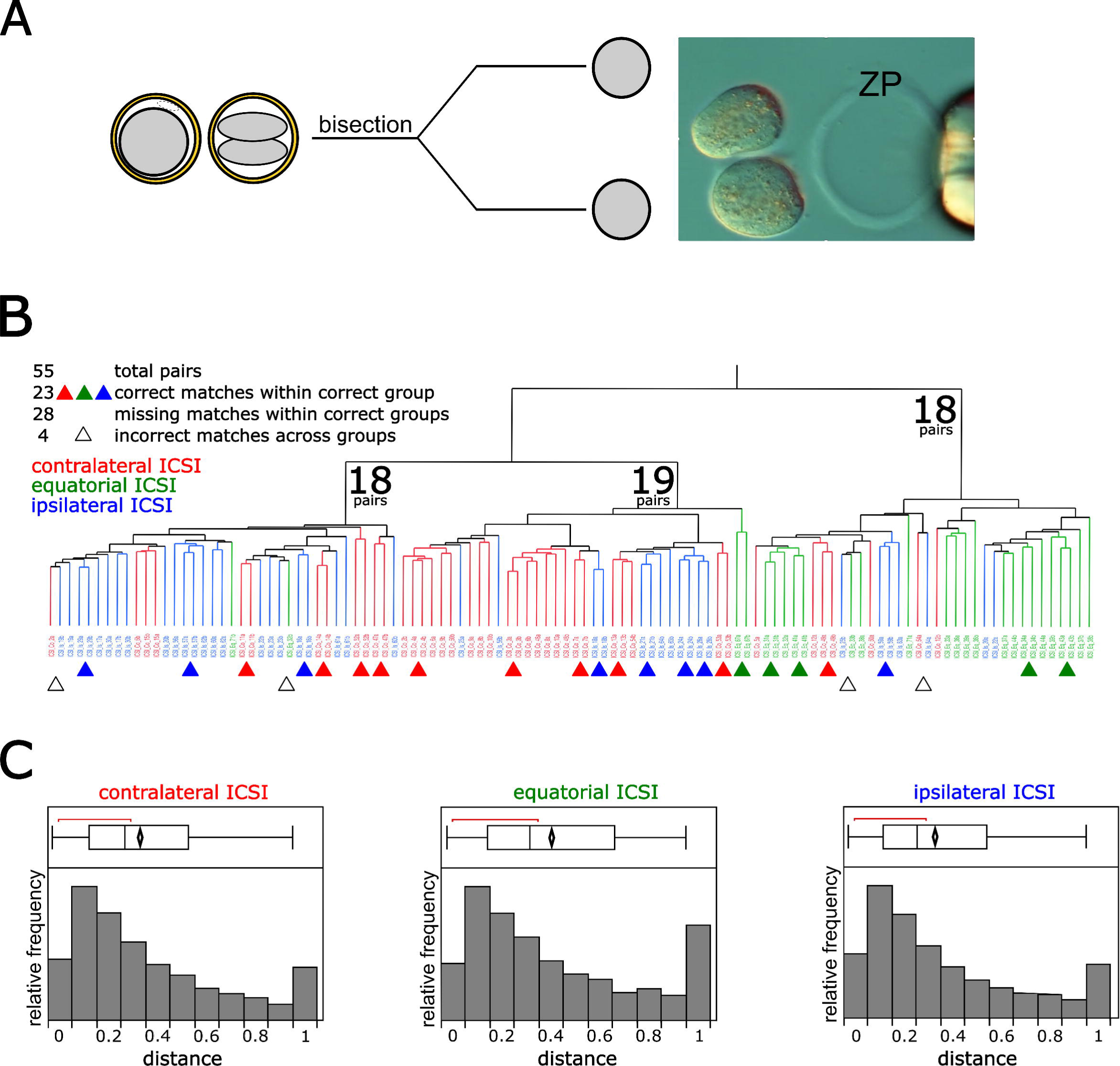
Single-cell RNA-seq of sister blastomeres separated after site-specific ICSI. **(A)** Scheme and representative image of a 2-cell embryo splitting. **(B).** Non-supervised hierarchical clustering analysis (method of Ward) of individual blastomeres’ transcriptomes (5 5 pairs, 7492 mRNAs). The coding system is as follows: ICSl_’Co’ or ‘Eq’ or ‘Is’ _Embryo number_’a’ or ‘b’, to indicate the one or the other blastomere (’a’ or ‘b’) of a given embryo (’Embryo number’) from contralateral, equatorial or ipsilateral ICSI. One can see from the colors that the blastomeres reflect the ICSI groups, and that most blastomeres are paired correctly with the sister blastomere. **C).** Distributions of inter-blastomere distances (ratio TPM difference/ TPM sum) after contralateral, equatorial and ipsilateral ICSI. Values of ‘0’ and ‘l’, forexample, indicate that the sister blastomeres have maximum reciprocal similarity and maximum reciprocal diversity, respectively, for the genes concerned. One can see that the histogram distributions of contralateral and ipsilateral ICSI are indistinguishable in spite of the opposite fertilization topologies (contralateral vs. ipsilateral ICSI, p = 0.311; Wilcoxon test), while either differs from the equatorial distribution (p < 0.0001; Wilcoxon test). Abbreviations: ZP, zona pellucida.

In addition to the technical quality control (QC) of the RNA-seq pipeline, we also filtered these mRNAs so as to retain those that had a gene name assigned and that were expressed consistently above background (see Supplementary Table S1 for more information on the criteria applied to include genes in the final analysis). By these criteria, 7492 mRNAs were retained across 21, 13 and 21 pairs of 2-cell stage blastomeres fertilized contralaterally, equatorially and ipsilaterally, respectively (55 pairs = 110 blastomeres in total). Gene expression levels were rendered as transcripts per million (TPM). The higher QC rejection incurred by the results of the equatorial group remains unexplained. A summary table of the 7492 mRNAs and their TPMs in each of the 110 blastomere transcriptomes is provided in Supplementary Table S2. The raw RNA-seq data are deposited in Gene Expression Omnibus under accession number GSE241089.

We subjected the 110 individual transcriptomes to non-supervised hierarchical clustering, expecting it to return the ICSI groupings (contralateral, ipsilateral, equatorial ICSI) and the native pair associations. Indeed, the clustering reflected the original groups and paired the blastomeres correctly with few gross mis-classifications (Figure 3B). We then tested a hypothesis that the three fertilization topologies would leave distinct footprints in the global gene expression, whereby inter-blastomere differences would be more or less pronounced depending on the ICSI group. We faced a problem that it is not, *a priori*, possible to know which blastomere in a pair is “first” and which blastomere is “second” so as to perform ratios or subtractions of transcript levels in a consistent manner across all 110 pairs of blastomeres. While we could know it *a posteriori* based on e.g. which blastomeres undergoes first the next mitosis, the caveat is that it would no longer be a 2-cell stage blastomere. To overcome this problem, we devised a simple analytical strategy, as follows. For each pair of blastomeres, we calculated the absolute difference of the two TPMs and divided it by the sum of the two TPMs, for each gene. This calculation not only is free from arbitrary decisions such as deciding which blastomeres to put in the numerator and which in the denominator (both are featured in the numerator as well as in the denominator), but it also provides a normalization (the range is between 0 and 1). After this calculation, a value of ‘0’ means that sister blastomeres have the same TPMs, whereas a value of ‘1’ means that the mRNA is detected in one blastomere but not in the other. Over all samples of each group, we called this adimensional ratio the ‘distance’ of the sister blastomeres from each other in terms of gene expression. A summary table of the 55 inter-blastomere distances is provided in Supplementary Table S3. The plots of the inter-blastomere distances for all 7492 genes returned three distributions that were highly similar and, in case of the contralateral and ipsilateral ICSI, also statistically indistinguishable (Figure 3C). This is not consistent with our hypothesis that the three fertilization topologies would leave distinct footprints in the global gene expression.

We entertained a possibility that the inter-blastomere differences might have been latent at 1^st^ division, but emerged later as the blastomeres progressed to blastocysts. We therefore separated and isolated the sister blastomeres from a new set of ICSI zygotes (experimental settings unchanged), but instead of lysing the blastomeres, we cultured them apart while keeping track of the original pair associations. This allowed us to generate twin and cotwin blastocysts (Figure 4A) as per our established method (Casser *et al*., 2017). Fourteen pairs of individual blastocysts from each ICSI group (total n=42) were processed for RNA-seq, with an average sequencing depth 2.9 ± 1.1 million uniquely mapped reads per specimen of single blastocysts. After applying same pipeline and same QC criteria as for the blastomeres (Supplementary Table S1), 7660 mRNAs were retained across 6, 7 and 6 pairs (total pairs n=19) of twin blastocysts from the contralateral, ipsilateral and equatorial group, respectively (GSE241089, Supplementary Table S4). As done for the blastomeres, we subjected the 38 transcriptomes to non-supervised hierarchical clustering (Figure 4B) and reduced the dimensionality of the data by calculating the ‘distances’ between twin and cotwin blastocyst (ratio TPM difference / TPM sum) for each gene, averaged over all samples (Supplementary Table S5). Unlike the blastomeres, the clustering of blastocysts did not reflect the original groups (Figure 4B); but like the blastomere it is evident how similar the inter-blastocyst distances are, even though the statistical test yields a significant p value (Figure 4C). Again, this is not consistent with our hypothesis that the three fertilization topologies would leave distinct footprints in the global gene expression.

**Figure 4.**
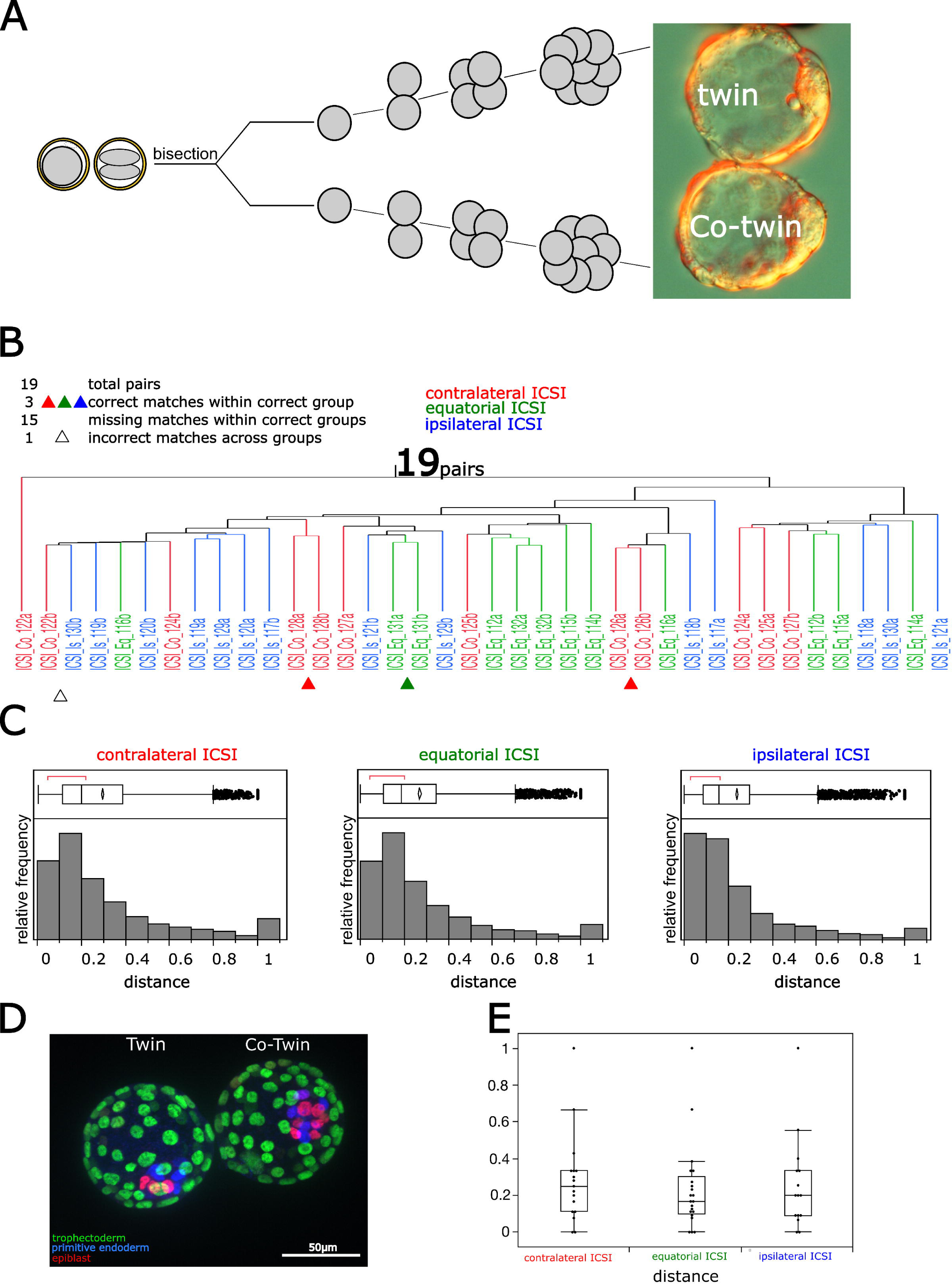
Single-embryo RNA-seq of twin blastocysts formed after site-specific ICSI. **(A).** Scheme and representative image of twin blastocysts from a split 2-cell embryo. **(B).** Non­ supervised hierarchical clustering analysis (method of Ward) of individual blastocysts’ transcriptomes (19 pairs, 7660 mRNAs). The coding system is as follows: ICSl_’Co’ or ‘Eq’ or ‘Is’ _Embryo number_’a’ or ‘b’, to indicate the one or the other twin (’a’ or ‘b’) of a given embryo (’Embryo number’) from contralateral, equatorial or ipsilateral ICSI. One can see from the colors that the twin blastocysts no longer reflect the ICSI groups. **C).** Distributions of inter-blastocyst distances (ratio TPM difference / TPM sum) after contralateral, equatorial and ipsilateral ICSI. Values of ‘O’ and ‘1’, for example, indicate that the twin blastocysts have maximum reciprocal similarity and maximum reciprocal diversity, respectively, for the genes concerned. One can see that the histogram distributions are highly similar, although their reciprocal comparisons yield a significant p value (p < 0.0001; Wilcoxon test). **(D)** Representative immunostaining of twin blastocysts for protein markers of epiblast (Nanog, red), primitive endoderm (Sox17, blue) and trophectoderm (Cdx2, green). (E) Distribution of inter-blastocyst distances based on the counts of epiblast cells (p > 0.361; Wilcoxon test). N of scored blastocysts: contralateral ICSI = 16; equatorial ICSI =20; ipsilateral ICSI = 14.

As a last possibility, we recalled from our original study (Casser *et al*., 2017) that the differences between twin and cotwin blastocysts were not as pronounced at the RNA level as they were at the protein level, particularly in the protein markers of the germ layers (trophectoderm, primitive endoderm, epiblast). This prompted us to apply germ layer immunofluorescence also in the case of the present study. Specifically, we applied Cdx2 (trophectoderm), Sox17 (primitive endoderm) and Nanog (epiblast) immunofluorescence on a new set of twin blastocysts obtained after contralateral, equatorial and ipsilateral ICSI (Figure 4D). In particular, we were interested in the epiblast, as this germ layer was the one with the most pronounced imbalance between twin and cotwin (Casser *et al*., 2017). As done for the transcriptomes we converted the data into adimensional distances, by calculating the absolute difference of cell counts in the two epiblasts and dividing it by the sum of cell counts, for each pair of twin blastocysts (Supplementary Table S6). In line with the distances in mRNA expression, also the statistical distributions of epiblast distances were similar to each other (Figure 4E). Thus, also the immunofluorescence assay was in line with the RNA-seq results leaning toward a refutation of our hypothesis.

The results of this section can be summarized as follows. In spite of the standardization enabled by imposing 3 fertilization sites (contralateral, equatorial, ipsilateral), the sister blastomeres differed from each other in the same way, and so did also the twin blastocysts. The categorization of the fertilization site into three classes (contralateral, equatorial, ipsilateral) did not result in three classes of embryonic products, but in one.

### Effect of oocyte geometry on the partition of oocyte territories is stronger than that of fertilization site

The results of blastomere and blastocyst analyses suggested that either the oocyte territories did not matter, or something prevented them from making a difference. The latter could be a meridional division, whereby zygotes receive equal portions of animal and vegetal materials, in spite of three different fertilization topologies. In dealing with these possibilities, we faced a problem that oocytes roll with the ZP and rotate within the ZP (dubbed ‘acrobatics’; (Graham *et al*., 2021)). Therefore, we secured an invariant view of the angle subtended between A-V axis of oocytes and cleavage axis of zygotes, by holding the sperm-injected oocytes physically immobilized one at a time in the micromanipulation chamber for 24 h (Figure 5A,B). For documentation, still pictures were preferred over time-lapse, in order to minimize phototoxicity, and pictures were taken at three time points i.e. at the time of ICSI as well as 5 and 24 hours later (Figure 5C). The unchanged position of the ICSI hole in the ZP and the dent left by the ICSI needle in the oolemma (examples shown in Figure 5C, white arrows and circles) attest that the immobilization of the oocytes was successful.

**Figure 5.**
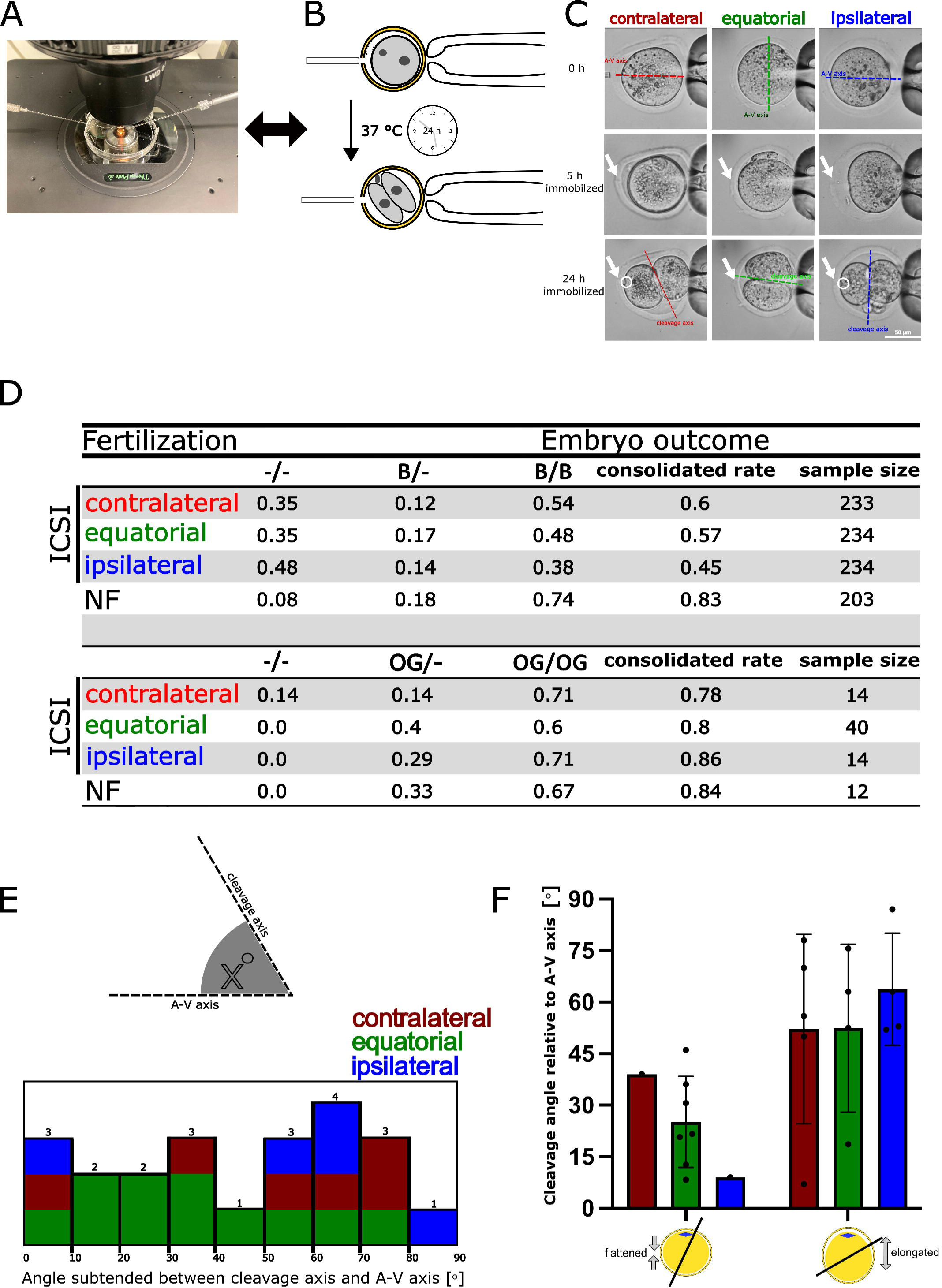
Biological lessons learned from the immobilization of oocytes for 24 h after site-specific_°_ ICSI. **A.** Overview of the micromanipulation chamber placed on Thermo Plate at 37 Con the stage of the inverted microscope. B. One oocyte (B6C3Fl) at a time was injected with a single sperm head (CDl). After injection the oocyte was not released, but held in position (immobilized) for 24 h in the micromanipulation chamber, thereby allowing for pronuclear formation and 1^st^ cleavage. **(C).** Representative images of oocytes of each ICSI group, showing the A-V axis of oocytes, the cleavage axis, and the ICSI hole in the ZP (white arrow) as well as the dent left by the ICSI needle in the oolemma (white circle). **(D)** Developmental validation of ICSI zygote viability after 24h immobilization. The sister blastomeres of 2-cell embryos were mechanically separated from each other. After splitting, they were followed up through 72 h of development post-bisection i.e. approx. 96 h from ICSI. Given the paired nature of the data, blastocyst (B) formation was scored as B/B (observed in both members of the pair), B/-(observed in one member of the pair), or -/-(observed in neither member). The consolidated rates (1x rate B/B + ½ x rate B/-+ 0 x rate -/-) were different among the ICSI groups (p = 0.0044, chi-square test) and were also lower for ICSI than for NF (p = 8.lE-13, chi-square test). Blastocysts were assayed for further development via the outgrowth (OG) assay. Also in this case, given the paired nature of the data, OG formation was scored as OG/OG (observed in both members of the pair), OG/-(observed in one member of the pair), or -/-(observed in neither member). Total N of pairs examined are provided as ‘sample size’. **(E).** Distribution of angles (°) subtended between A-V axis and 1^st^ cleavage axis. Note that while the ipsilater_°_al and contralateral ICSI populated the high end of the angle distribution (60-90°), the equatorial ICSI was distributed more evenly across angles. (F). Distribution of angles (°) presented in (E), as a function of the oocyte being flattened vs. elongated in the direction of the A-V axis. Abbreviations: B, blastocyst; NF, natural fertilization; OG, outgrowth.

We ascertained that the immobilization of oocytes during ICSI and for 24h thereafter did not compromise the developmental ability of the zygotes. Given that only a single 2-cell embryo could be produced per day due to the immobilization on stage for 24h, embryo transfers to pseudo-pregnant females were not feasible. Therefore, we applied *in vitro* functional assays to test if the 2-cell embryos could form blastocysts (in incubator) when either preserved intact or separated into the two blastomeres. In order to test the immobilized 2-cell embryos we extracted both blastomeres one after the other through a slit made in the ZP, as per our established method (Casser *et al*., 2017). Only pairs in which both of the blastomeres survived were followed up, since it is not possible to study the partition of pole materials in the daughter cells, unless they both survive (Figure 5D). Rates of dual blastomere survival during bisection were similar across the three groups (contralateral ICSI; 64% ± 12%, 7 sessions, 233 pairs survived; equatorial ICSI: 60% ± 18%, 7 sessions, 234 pairs survived; ipsilateral ICSI: 68% ± 16%, 7 sessions, 234 pairs survived) albeit lower than those of naturally fertilized counterparts (83 ± 16%, 6 sessions, 203 pairs survived). We speculate that embryos obtained by ICSI are more fragile during the splitting.

After four days of culture as per our established method (Casser *et al*., 2017), the rates of blastocyst formation were recorded and compared with those of intact counterparts (38-43%, Figure 2A). We faced an issue that since one bisected 2-cell embryo becomes two embryos, there can be three outcomes: two blastocysts (B/B), only one blastocyst (B/-), or none (-/-). To compare this multiplicity with the single outcome of the intact counterparts, the three outcomes were consolidated (1 x rate B/B + ½ x rate B/-+ 0 x rate -/-). As already noted, the reduction in blastocyst formation was probably the tradeoff for the higher consistency of using the same batch of cryopreserved spermatozoa throughout the study. The twin blastocysts (B/B) were tested further using an *in vitro* model of implantation known as the ‘outgrowth assay’ (Armant *et al*., 1986; Kim *et al*., 2020), in which the blastocysts are deposited onto a layer of mitotically inactivated fibroblasts to see if they attach and their trophectoderm proliferates (grows out). Unlike the difference in blastocyst formation, no major differences were seen among the groups tested for outgrowth formation (Figure 5D) (p>0.04, chi-square test).

Given that immobilization and imaging preserved viability, we assumed that the events observed here *in vitro* reflected those *in vivo*, and used the pictures to draw biological lessons. We analyzed the angle subtended between cleavage axis and A-V axis of 22 individual oocytes (Figure 5E). The angle was autonomous to the oocyte: when the oocyte was rotated by 90° (all other parameters unchanged), the cleavage axis changed too, following the new orientation of the oocyte, as exemplified in the middle column in Figure 5C. Taking all the 22 cases, however, we realized that the angles did not relate in any simple way to mainstream models of 1st cleavage in mice (see the Introduction). Whereas one model predicates that the first cleavage occurs parallel to the A-V axis or within 30° of it, which is referred to as ‘meridional’, we observed pure meridional cleavage only in a minority of cases (n = 3/21; left-most bar in Figure 5E). Whereas the other model predicates that cleavage is perpendicular to the A-V axis i.e. equatorial only when the sperm enters the oocyte at the vegetal pole, perpendicular cleavage occurred also after ICSI at the animal pole. We took a closer look at the cases of equatorial cleavage (60-90° in Figure 5E) and broke them down by ICSI type: 37% resulted from contralateral ICSI, 37% from ipsilateral ICSI, 26% from equatorial ICSI. In other words, zygotic division near the equator was prevalent (74%) when the fertilization site laid on the A-V axis of the oocyte, which is a prevalence strikingly similar to that of another study (Zheng *et al*., 2013) in which the frequency of equatorial cleavage was quantified in 73%. It was while the images were being examined that also a new observation related to the oocyte geometry was made. We noted that that the shape of the oocytes was oval rather than spherical, thus having a short and a long diameter, and that the A-V axis of oocytes was often – albeit not always - the longest. This reminded us of the Hertwig’s rule (Hertwig, 1893), whereby cells tend to divide across the shorter diameter. This prompted us to reexamine the image records (Figure 5C). After scoring the 22 images again, we realized that when the MII oocyte appeared flattened in the direction of the A-V axis prior to ICSI, the oocyte tended to divide along that axis (meridional) after ICSI; whereas when the oocyte appeared elongated in the direction of the A-V axis prior to ICSI, the oocyte tended to divide perpendicular to that axis (equatorial) after ICSI (Figure 5F).

The results of this section can be summarized as follows. Mouse oocytes had a strong preference to divide across their short diameter, irrespective of the fertilization site, apparently following the Hertwig’s rule. This way of division explains why the imposition of fertilization at three different locations by ICSI resulted in one class of 2-cell embryos rather than three (the shorter diameter is one), and why – on average - the sister blastomeres differed from each other always in the same way (the animal and vegetal materials segregated in majority of 2-cell embryos irrespective of which ICSI group). Our methodological approach reveals that the contribution of ooplasmic territories to the variability of blastomeres is constrained by a mode of cell division – along the equator - hitherto regarded atypical for mouse oocytes.

## Discussion

In order to test if it is the orientation of 1^st^ zygotic division what makes the blastomeres of 2-cell embryos differ from each other (Seidel, 1952; Seidel, 1960) and accounts for the observation that monozygotic twin embryos are different (Casser *et al*., 2017), we standardized the topology of fertilization *in vitro*. We harnessed the ICSI method (Kimura and Yanagimachi, 1995) to deposit the sperm head at the animal pole, vegetal pole or equator of mouse oocytes. We call it “site-specific ICSI”. If oocytes had instead been fertilized *in vivo* or inseminated *in vitro*, the fertilization site would have been random albeit with a preference for the oocyte’s equator and an aversion to the animal pole (Mori *et al*., 2021; Piotrowska and Zernicka-Goetz, 2001; Piotrowska-Nitsche and Chan, 2013). This would have amounted to conducting a study with a mixed population in which each zygote was different from the other, yielding unreproducible results. Instead of this natural variability, our ICSI zygotes reflected only three – specific and reproducible - sites of fertilization. Although the jury is still out in regards to mouse ICSI embryos being equivalent (Piotrowska-Nitsche and Chan, 2013) or not equivalent (Giritharan *et al*., 2010) to natural counterparts, the differences are probably not in fundamental biological properties, as suggested by the millions of ICSI babies born in medically assisted reproduction worldwide. In our ICSI setting all three fertilization topologies were developmentally successful, including that at the animal pole, which is usually recommended to avoid in ICSI because of the risk of perturbing the meiotic spindle (Blake *et al*., 2000; Labs *et al*., 2016; Van Der Westerlaken *et al*., 1999).

Based on current knowledge (Introduction), cleavage axes orient relative to a multiplicity of cues that have different strengths and competitive hierarchal relationships with one another. While the exact outcomes are difficult to predict, the three fertilization sites of this study were expected to result in different orientations of the zygotic cleavage plane. Therefore, if we accept that the animal and vegetal hemispheres differ in composition (former is enriched in spindle-associated transcripts (VerMilyea *et al*., 2011), latter is richer in endoplasmic reticulum (FitzHarris *et al*., 2003; Kline *et al*., 1999)), then the different orientations of 1^st^ cleavage should result in different partitions of animal and vegetal materials into the blastomeres of the 2-cell stage embryo. This could be a simple cytological basis to explain the non-balanced properties of monozygotic twin blastocysts produced by splitting of 2-cell embryos in mice (Casser *et al*., 2017). Therefore, we compared and contrasted the sister blastomeres of our 2-cell embryos obtained after contralateral, equatorial and ipsilateral ICSI, using single-cell RNA-seq. This is not the first time that individual sister blastomeres of 2-cell mouse embryos have been subjected to transcriptome analysis (Biase *et al*., 2014; Piras *et al*., 2014; Shi *et al*., 2015; VerMilyea *et al*., 2011). However, it is probably the first time that sister blastomeres were produced in a way that minimized the fertilization variability and were analyzed in a way that minimized batch effects through the use of Mosquito® plate. On the day following site-specific ICSI, sister blastomeres were dissociated and subjected to single cell RNA-seq or cultured to blastocyst stage followed by single embryo RNA-seq. We reduced the dimensionality of the transcriptomic data, by calculating the adimensional ‘distances’ between the blastomeres (TPM difference / TPM sum). Using these distances, we generated statistical distributions (histograms) of the three ICSI groups, expecting them to be well distinguishable from each other or even to form three non-overlapping distributions. Instead, the three distributions turned out to be almost identical, as if to suggest that although ooplasmic territories of various composition may exist, their allocation to the blastomeres is functionally irrelevant. We reasoned that perhaps the 2-cell stage was still too early and latent differences would emerge during subsequent development. We separated the sister blastomeres and cultured them apart, in parallel, so as to produce twin blastocysts, as per our established method (Casser *et al*., 2017). These blastocysts were subjected to single embryo RNA-seq, using the same pipeline of the blastomeres. In addition, we recalled that that the twin blastocysts of our previous study (which was based on *in vivo* fertilization) presented systematic differences in the epiblast, which was bigger in one twin than in its cotwin (Casser *et al*., 2017). Therefore, we counted the cells in the germ layers of twin blastocysts resulted from site-specific ICSI and converted them into distances. Again, the three distributions of inter-blastocyst distances turned out to be almost identical, both at the level of transcripts and at the level of epiblast cell numbers. Either the oocyte territories did not matter, or their effect was evened out by other factors.

To shed light on the elusive factors, we introduced the physical immobilization of the oocytes during ICSI and for the 24h thereafter, in order to ascertain the transmission of animal and vegetal materials to the first two blastomeres in the absence of oocyte rolling and rotation. This way we realized that the oocytes fertilized by ICSI at the animal pole (ipsilateral) or vegetal pole (contralateral) had a strong bias (>70%) to divide perpendicular to the A-V axis. While this cleavage orientation is detrimental for development in sea urchin (Gardner, 1996), the jury is still out when it comes to mammals. Cytological evidence is limited to a handful of studies in which the occurrence of oocytes’ movements could not be ruled out completely. In one study, naturally fertilized mouse oocytes were reported to divide perpendicular to the A-V axis, which was marked at the microscope by way of two microdrops embedded in the ZP, one above the 2^nd^ PB and the other at the opposite site. On the assumption that the zygotes retained the initial position (i.e. did not rotate inside ZP) when moved to the incubator and then returned back to the microscope on the following day, it was concluded the 1^st^ division had occurred equatorially (Figure 3C,D and Figure 4b in (Gardner and Davies, 2003)). More recently, also another study reported that a substantial proportion (73%) of mouse zygotes divided equatorially (Zheng *et al*., 2013). Apart from mice, equatorial division was observed in ICSI zygotes of Rhesus monkeys (Simerly *et al*., 2019). An important difference between our study and the prior studies in mice is that we did not assume lack of oocyte rolling and rotation – we imposed it (our oocytes were immobilized using a micromanipulator). Under our standardized conditions we realized that the equatorial division was more common than we thought, particularly when oocytes were fertilized consistently at one pole or the opposite pole by ICSI. Unlike these oocytes, those we injected at the equator appeared to have more variability in the angle subtended between 1^st^ cleavage axis and A-V axis. This variability is reminiscent of observations made by other scientists after polar material was removed from mouse zygote by cuts made in different position along the A-V axis. Quote: “cutting eggs perpendicularly to the AV axis caused more of them (56%) to change the cleavage plane than cutting meridionally (29%)” (Zernicka-Goetz, 1998), suggestive to the Authors of factors distributed along the equatorial region with roles in the orientation of the cleavage axis. We too observed a variability associated with the equatorial region, but it turned out to be apparent. When examining the pictures, we serendipitously noted that the A-V axis of oocytes was consistently the longest, albeit with exceptions. This reminded us of the Hertwig’s rule (Hertwig, 1893), whereby cells tend to divide perpendicular to their long axis. Indeed, it was already known that when mouse zygotes were compressed laterally, thereby elongating the A-V axis, they tended to cleave along their newly acquired shorter diameter (Gray *et al*., 2004). We envisioned that similar factors might have played a role also in our ICSI study. Indeed, upon re-examination of our data, we found that regardless of the fertilization site, the 1^st^ zygotic division was strongly associated with the shorter diameter of the MII oocyte. Thus, the Hertwig’s rule could in itself already explain why there was a bias toward equatorial division after ICSI conducted at the animal or vegetal pole. A limitation of our study is that we did not examine the oocytes and zygotes for intra-ooplasmic movements (Ajduk *et al*., 2011; Graham *et al*., 2021). These may have well occurred and accounted for the residual variability i.e. the standard deviation of the angles subtended between A-V axis and 1^st^ cleavage axis. We also did not examine shape changes and flattening of oocytes occurring after fertilization (Gray *et al*., 2004). Our oocytes were immobilized, and either with or without changes possibly occurring after fertilization, fact remains that the pre-fertilization short diameter already predicted how the oocyte would divide after fertilization.

In conclusion, this study reported an original approach for removing some of the confounders that affect the study of inter-blastomere differences and blastomere diversification at the beginning of mouse development. Our results support that even if we standardize the topology of fertilization, the oocyte is not perfectly round, and decisive is the shorter diameter. As the A-V axis is typically the longer diameter, this has as consequence that the oocytes divide equatorially, bringing about a segregation of animal and vegetal materials in the 2-cell stage blastomeres. This offers a rationale to explain the distinct properties of sister blastomeres and resultant monozygotic twins, since one receives more of the animal materials and the other more of the vegetal materials. Our micromanipulation combined with immobilization provides an opportunity for further interrogation of the roles of oocyte territories, toward a more complete understanding of their contribution in the genesis of inter-blastomere differences in mice.

## Materials and methods

### Compliance with regulations on research animals

All mice used were maintained in individually ventilated cages in the animal facility of the MPI Münster, with a controlled temperature of 22 °C, a 14/10 h light/dark photoperiod and free access to water and food (Harlan Teklad 2020SX). Procedures used in this study followed the ethical guidelines of the European Laboratory Animal Science Associations (FELASA) and the ARRIVE reporting guidelines. On the local regulatory level, mice were used for experiments according to the ethical approval issued by the Landesamt für Natur, Umwelt und Verbraucherschutz (LANUV) of the state of North Rhine-Westphalia, Germany (Permit number 81-02.04.2020.A405, “Die Zuteilung der zygotischen Totipotenz in den ersten Blastomeren: Eine Untersuchung der Ursprünge und Mechanismen im Mausmodell”).

### Collection of oocytes

Six-to eight-week-old B6C3F1 females were primed with 10 I.U. each of pregnant mare serum gonadotropin (PMSG, Intergonan, Intervet) and human chorionic gonadotropin (hCG, Ovogest, Intervet) injected intraperitoneally 48 h apart at 5pm, and then killed by cervical dislocation to collect MII oocytes. Cumulus cells were removed in hyaluronidase (50 I.U./mL in HEPES-buffered CZB medium; (Cavaleri *et al*., 2006)).

### Collection of sperm and cryopreservation

Caudae epididymis from two 3-month-old CD1 male mice were collected, and spermatozoa were allowed to swim-up in ∼ 1 mL Whittingham medium (supplemented with 3% BSA, fraction V) across a distance of ∼ 1.5 cm for 30’ under a humidified atmosphere of 5% CO_2_ in air. The upper layer (∼ 200 μL) of medium was collected and incubated under pre-equilibrated mineral oil, to allow for sperm capacitation. After 1h the drop of medium containing the spermatozoa was centrifuged and then resuspended in half the volume of Whittingham medium. Aliquots of 5 µL of sperm suspension were placed at −80 °C for one week and then stored in liquid nitrogen.

### Intracytoplasmic sperm injection (ICSI) and embryo culture

MII oocytes were subjected to ICSI in HEPES-buffered CZB medium as per our micromanipulation protocol in which the holding pipette is on the right-hand side and the ICSI pipette comes from the left, except that this time they were also rotated to consistently inject the sperm head at 03:00 with the MII spindle at 09:00 (vegetal pole ICSI), at 03:00 (animal pole ICSI) or at 12:00 (equatorial ICSI). When applicable, the oocytes were also immobilized. A slight but constant suction was applied all the time with the holding pipette. To assess the relationship between site of ICSI and 1^st^ zygotic cleavage, we kept the microinjected oocytes on the stage of the micromanipulator. We empirically found that cleavage was supported on plastic bottom (no glass) under air, by adding the micromanipulation medium with 25% of KSOM(aa) and preserving osmolarity via oil overlay, at a temperature of 37 °C maintained via Thermo Plate (Tokai Hit). Once the relationship was determined, in the subsequent experiments the injected oocytes were transferred to 500 µL of KSOM(aa) embryo culture medium in a 4-well plate without oil overlay, at 37 °C under 6 % CO_2_ in air, until bisection on the next day.

### 2-cell embryo bisection

Two-cell embryos arising in the time frame from 6 to 7 am were removed from culture to perform splitting from 8 to 9 am. These embryos were transferred in groups of 12 to a micromanipulation drop on the stage of a Nikon Eclipse TE2000-U inverted microscope fitted with Nomarski optics and holding and bisection needles in place. The micromanipulation medium consisted of 0.2 mM D(+) glucose, 0.2 mM pyruvate, 10 mM lactate, 0.5% w/v BSA, in 0.9% w/v sodium chloride, in place of the commonly used calcium-magnesium-free phosphate-buffered saline whose phosphate was shown to be detrimental to embryo development (Scott and Whittingham, 1996). We also made the bisection medium slightly hypertonic by using 95% of the water volume. The bisection needle was a TransferTip needle operated by a CellTram Vario (Eppendorf). The two-cell embryo was rotated using the holding and bisection needle to align the cleavage plane with the common axis of the two pipettes. The 2-cell embryo was firmly held in place with the holding pipette by applying negative pressure (suction). The TransferTip was used to first make a slit in the ZP at one pole and then to press the ZP gently at the equatorial region, causing one blastomere to be squeezed out. Suction in the holding pipette was reduced. The other blastomere came out by performing the same procedure, except that the ZP was pressed below the equatorial region. When all 12 embryos had been bisected, which normally took 8–10 min, the pairs of blastomeres were collected and transferred to the 96-well plate (round bottom) using a bent (≈60° angle) mouth-operated pipette with flame-polished tip. The individual blastomeres were either lysed in RNA buffer or allocated individually to embryo culture in a 96-well plate with round bottom (75 µL medium per well, without oil overlay). Culture medium was KSOM(aa). Embryos were cultured for an additional 72 h at 37 °C under 6 % CO2 in air until the blastocyst stage.

### Embryo transfer (ET) and post-implantation development

Blastocysts from intact or bisected 2-cell embryos (72 h after bisection) were transferred either as pools of eight or as single pairs (twin and co-twin, with six oocytes as carriers) to one uterine horn of pseudo-pregnant CD1 recipients that had the copulation plug from vasectomized CD1 males three days prior to the ET. Typically, the recipients weighed between 25 and 30 grams and were older than 6 weeks but no older than 3 months. Prior to surgery, CD1 recipients were anesthetized with Ketamine (80 mg/kg)/Xylazine (16 mg/kg)/Tramadol (15 mg/kg) in PBS, delivered intraperitoneally to the mouse at 10 μl/g body weight. Embryos were delivered to the uterine lumen using a mouth-operated glass pipette through a hole made in the cranial region of the uterine horn using a 27G needle. Wounds in the skin were closed with metal clips. Typically, the surgery per se took 10-15 minutes per mouse. The post-surgery recovery area was warmed to 30 °C using infrared lamps. Pregnancies were recorded by cesarean section shortly prior to the natural term (embryonic day (E) 18.5).

### In vitro model of implantation (outgrowths)

Using our adaptation (Boiani *et al*., 2002) of the outgrowth method (Armant *et al*., 1986) we transferred twin blastocysts (72 h after two-cell embryo bisection) individually onto a feeder layer of γ-ray-inactivated mouse embryonic fibroblasts (C3H background) grown to confluence in 96-well plates (flat bottom) previously. The medium consisted of high-glucose DMEM (Gibco) with 15% fetal bovine serum (BioWest, Nuaillé, France), glutamine and penicillin/streptomycin (Gibco), non-essential amino acids (PAA Laboratories, Pasching, Austria), mercaptoethanol 0.1 mM (Gibco), 1000 Units/mL LIF (produced in-house).

### Transcriptome analysis of single blastomeres and single blastocysts

Single blastomeres or single blastocysts were lysed in 10X Lysis Buffer from Takara (cat.no. 635013), deposited in two Mosquito® plates, for a total of 150 pairs of blastomeres (300 cells) and 78 blastocysts (36 intact, 42 twins from 21 pairs). Then the RNA was extracted and converted to cDNA using TaKaRa SMART-Seq Single Cell Kit (cat.no. 634470). Sequencing libraries were prepared using the Illumina Nextera XT DNA Library Preparation Kit (cat.no. FC-131-1024). Libraries were sequenced on an Illumina NovaSeq 6000 platform to obtain two datasets of 4.2 ± 1.5 million total mapped reads per library for blastomeres (dataset 1 with 150-base paired-end reads and dataset 2 with 100-base single-end reads). For blastocysts, the datasets had 2.9 ± 1.1 million total mapped reads per library (only 100-base single-end reads). In order to mitigate any potential bias arising from the discrepancies between these libraries, only the initial reads from the paired-end dataset were preserved, while the last 50 bp of each read were trimmed. Consequently, the subsequent blastomeres analysis exclusively employed sequencing data from both libraries, wherein the preserved first reads from the paired-end dataset were adjusted to a standardized length of 100 bp.

Trimmomatic V.039 (https://doi.org/10.1093%2Fbioinformatics%2Fbtu170) was applied to trim reads and remove adapters with the following parameters; “SE ILLUMINACLIP:NexteraPE-PE.fa:2:30:10 LEADING:3 TRAILING:3 SLIDINGWINDOW:4:15 MINLEN:50”. A pre-built mouse reference set was downloaded from the 10X genomics repository (https://cf.10xgenomics.com/supp/cell-exp/refdata-gex-mm10-2020-A.tar.gz) on 12.11.2022, which included the reference genome (GRCm38) and gene annotation (M23, Ensembl 98). To quantify gene expression, the reads originated from blastomeres and blastocysts were separately mapped to the mouse reference genome using the STAR pipeline (https://doi.org/10.1093/bioinformatics/bts635) with the following options; “--soloType SmartSeq--readFilesManifest manifest.tsv--soloUMIdedup Exact NoDedup -- soloStrand Unstranded--genomeDir path_to/enome_dir/--soloFeatures Gene SJ--outSAMtype BAM SortedByCoordinate--quantMode GeneCounts TranscriptomeSAM -- outTmpKeep All”. Of the starting 300 cells (150 blastomere pairs), 124 cells (62 pairs) were retained. Of the starting 78 blastocysts, 69 were retained.

The resultant raw count matrix derived from blastomeres, composed of 124 cells (62 twin blastomeres and 32285 genes), was processed using the SCANPY toolkit (Wolf *et al*., 2018). This phase involved generating QC metrics and eliminating cells and genes of poor quality. During this step, cells exhibiting fewer than 9500 detected genes and mitochondria gene counts exceeding 0.02 percent were excluded from subsequent analysis. Due to this filtering process, the number of cells decreased from 124 to 117, resulting in the loss of corresponding mates for some twin blastomeres. To address this, the unpaired cells were removed, leaving us with a final set of 110 cells (55 pairs) for further analysis. Subsequently, those genes that were expressed in less than two cells and had fewer than ten reads mapped to them were filtered out. These filtering steps reduced the initial 32285 genes to 21778. Afterward, each gene’s transcript per million (TPM) was calculated using an in-house Python script. To accomplish this, the length of each gene was obtained utilizing the GenomicFeatures package (Lawrence *et al*., 2013). Following that, the median TPM value was calculated for each gene across 110 blastomeres. The genes were then sorted in descending order based on their median TPM values. In a subsequent step, genes annotated as ‘predicted genes’ were removed from the dataset. Finally, only those genes with a non-zero median TPM were retained, resulting in a total of 7492 genes for further analysis. The filtering steps are summarized in Supplementary Table S1.

The analysis of the count matrix corresponding to single blastocysts (dataset 2) followed the blastomeres procedure, with a few distinctions in the cell filtering criteria. Cells characterized by fewer than 4000 detected genes were excluded, and unlike blastomeres, no filtering was applied based on detected mitochondrial genes due to their normal counts. The subsequent steps in blastocyst processing, including TPM calculation, mirrored those applied to blastomeres as well. Consequently, the processed blastocysts encompassed a total of 69 specimens, comprising 38 twin blastocysts (19 pairs) and 31 intact blastocysts. Intact blastocysts were defined as those in which the 2-cell embryo was not split (unlike the twins). This dataset, comprising 7660 genes, was subjected to further analysis. The filtering steps are summarized in Supplementary Table S1.

### Analysis of cell lineage allocation in blastocysts

Blastocysts from intact or bisected 2-cell embryos (72 h after bisection) were analyzed by performing an immunostaining followed by confocal microscopy imaging to identify and map the different cell lineages, as described (Casser *et al*., 2017; Schwarzer *et al*., 2012). The following primary antibodies were applied simultaneously to the specimens overnight at 4 °C: Anti-Cdx2 mouse IgG1κ (Emergo Europe, The Hague, Netherlands, cat. no. CDX2-88), anti-Nanog rabbit IgG (Cosmo Bio, Tokyo, Japan, cat. no. REC-RCAB0002P-F) and anti-Sox17 goat IgG (R&D Systems, cat no. AF1924) in dilutions of 1:200, 1:2000 and 1:100, respectively. Appropriate Alexa Fluor-tagged secondary antibodies (Invitrogen) were matched to the primaries and incubated for 2 h at room temperature. Embryos were placed in 5 μL drops of PBS on a 50-mm thin-bottom plastic dish (Greiner Bio-One, Lumox hydrophilic dish; Frickenhausen, Germany) and overlaid with mineral oil (M8410 Sigma). Images were captured on the stage of an inverted microscope (Eclipse 2000-U; Nikon, Düsseldorf, Germany) fitted with a spinning disk confocal unit (Ultra View RS3; Perkin-Elmer LAS, Jügesheim, Germany). A Nikon Plan Fluor 40X oil immersion lens (NA 1.30) was used. Twenty optical sections per blastocyst were captured using a Hamamatsu ORCA ER digital camera (Hamamatsu Photonics KK, Japan). Maximum intensity projections were analyzed with ImageJ Version 1.46j, counting the CDX2-positive, SOX17-positive and NANOG-positive cells manually.

### Data and statistical analysis

The values presented in the figures are mean values ± standard deviations. Statistics was performed using Wilcoxon test or Chi-square test with JMP (SAS) software. Histograms and statistical distributions were generated in GraphPad Prism software.

## Data availability statement

All data needed to evaluate this article are provided in the main body or in the Supplementary Materials. Large-scale data (RNA-seq) have been deposited in the Gene Expression Omnibus of NCBI with accession number GSE241089. For convenience, the TPM of the RNA-seq datasets are summarized in the Supplementary Tables S2-S5 deposited in FigShare. The blastocyst cell counts and the inter-blastocyst distances presented in Figure 5 are provided in Supplementary Table S6 deposited in FigShare.

Supplementary Table S1: https://figshare.com/s/531f8d1b66fc523b2fea

Supplementary Table S2: https://figshare.com/s/9a68cf807f3c2ccf5c8a

Supplementary Table S3: https://figshare.com/s/4dcdcf44170061f69cde

Supplementary Table S4: https://figshare.com/s/b9c3993b82841b00699c

Supplementary Table S5: https://figshare.com/s/e05517c815dad487c920

Supplementary Table S6: https://figshare.com/s/9077fc19dacf44b774ac

## Acknowledgements

The authors would like to express gratitude for scientific environment and infrastructural support to the Max Planck Institute for Molecular Biomedicine. As part of infrastructural support, we experienced outstanding support from the mouse housing facility, ensuring a dependable supply of mice needed to collect oocytes and embryos. This study was in part presented orally to the ESHRE meeting 2023.

## Authors’ roles

According to CRediT (Contributor Roles Taxonomy; https://credit.niso.org) M.B., G.F. and W.M. conceived the study (<u>Conceptualization</u>) and acquired the funding (<u>Funding</u> <u>acquisition</u>). M.B. performed the micromanipulations, embryo culture, and embryo transfers (<u>Investigation</u>) and analyzed the results (<u>Formal Analysis</u>). S.I. purified the mRNAs for RNA-seq (<u>Investigation</u>). Y.S. provided infrastructure for the RNA-seq (<u>Resources</u>) and performed the RNA-seq (<u>Investigation</u>). R.H. performed the RNA-seq data analysis and curated the data under supervision of W.M (<u>Data curation</u>). T.N. and S.I. did the immunofluorescence imaging (<u>Investigation</u>). T.N. drew the figures (<u>Visualization</u>). M.B. wrote the initial draft (<u>Writing –</u> <u>original draft</u>). M.B., G.F. and W.M wrote the later versions of the manuscript up to its present form (<u>Writing – review & editing</u>).

